## Supplementary results for "Post-learning replay of hippocampal-striatal activity is biased by reward-prediction signals"

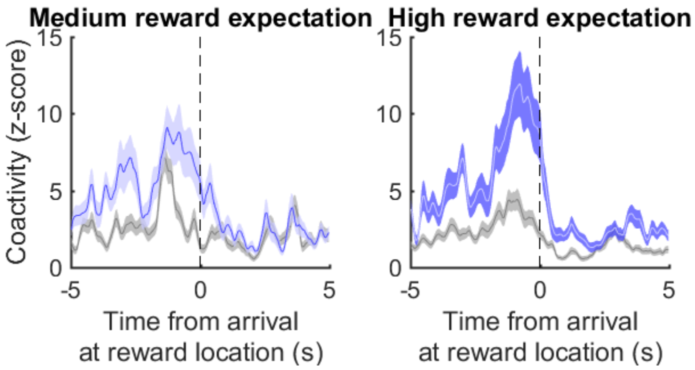


Figure S1: Mean ± s.e.m. z-scored coactivity of reactivated cell pairs (blue) and non-reactivated cell pairs (grey) around the time of arrival at reward locations on rewarded medium- and high-expected reward trials (rewarded outcomes only).
